# Microbially-mediated pretreatment of aerated oil and gas produced water by addition of phosphorous or activated sludge

**DOI:** 10.1101/2020.08.20.260448

**Authors:** Alexander S. Honeyman, Emily R. Nicholas, Tzahi Y. Cath, John R. Spear

## Abstract

Produced Water (PW) from oil and gas (O&G) producing wells is a unique source of water in water-stressed regions. Microbiota within O&G formations have been well-studied on site/*in-situ*, but few applied works have considered their role in the treatment of PW in engineered water treatment systems. Herein, we operated a simple aeration/mixing bench-scale bioreactor fed with produced water under three conditions: 1) PW alone (control; hereafter referred to as ‘baseline’), 2) phosphorous dosed daily as KH_2_PO_4_, and 3) activated sludge (AS) dosed daily from a sequence batch membrane bioreactor (SB-MBR). Aerated and mixed PW alone (baseline) was able to attenuate PW chemistry with removal of soluble chemical oxygen demand (sCOD) and ammonia by 27.6% and 17.8%, respectively. Further KH_2_PO_4_ and AS additions improved water treatment efficiency markedly; in the KH_2_PO_4_ addition reactor, sCOD and ammonia were reduced by 50.0% and 61.5%, respectively, and in the AS addition reactor by 52.5% and 59.2%, respectively. Microbial consortia determined via 16S rRNA gene amplicons differ in composition between raw PW and all reactors; order Kiloniellales was most common in raw PW while orders Rhodobacterales, Pseudomonadales, and Caulobacterales were most abundant amongst AS, KH_2_PO_4_, and baseline conditions, respectively. Several different microbial consortia are capable of treating raw PW which suggests that functional redundancy amongst microbiota in engineered treatment systems may be underappreciated. With simple addition of phosphorous and/or activated sludge to PW as part of a treatment strategy, a higher quality water can then be subjected to conventional treatment and/or local reuse.

**Importance:** Multiple microbiological communities are capable of treating O&G PW in a simple, applied, engineered setting. The broad possibility of PW treatment by multiple different microbial consortia elucidates the potential for easy, effective, water reuse processes in the hydraulically-stressed arid west as well as any region generating PW from O&G operations.

## 1. Introduction

Hydraulic fracturing of oil and gas (O&G) wells may induce additional water stress in regions already experiencing water shortages and drought (1). Hydraulic fracturing of deep formations is achieved by high-pressure injection of fluids and sand into horizontal wells, after which hydrocarbons are more easily released and brought to the surface. The recovered hydrocarbons are accompanied by water consisting of both injection fluids (also known as fracturing flowback fluids—FFBs) and formation fluids with varying concentrations of cations, anions, and organics. Combined, this fluid, broadly referred to as ‘produced water’ (PW) can be an important source of clean water if methods to properly treat it are developed. PW typically contains substantive concentrations of suspended solids, total dissolved solids (TDS), organic matter, volatile compounds, metals, total radioactivity (2), and microorganisms either native to the subsurface or introduced to PW by injection fluids (or both)—all of which complicate the treatment of PW by conventional water treatment processes. Thus, the treatment of PW for beneficial use is commonly substituted by injection to deep disposal wells or, otherwise, by non-beneficial use such as elimination and volume minimization in evaporation ponds (3). Removal of total organic carbon (TOC), suspended solids, total and soluble chemical oxygen demand (COD and sCOD), and ammonia would aid subsequent downstream treatment processes such as desalination.

Publicly owned treatment works (POTW) are not designed to handle the high salinity and concentration of metals / other dissolved inorganics native to PW (3). However, Frank et al. (4) show that treatment of PW in activated sludge (AS) systems is possible at loads of 6-20% PW by volume with limited impacts on the performance of the biological process and on effluent water quality. Microbial communities have also shown remarkable resilience to disturbance of batch reactor parameters during municipal water treatment (5), suggesting there may be an untapped potential in the capabilities of microbial communities for the treatment of PW. Pre-treatment of PW by microbial communities native to PW could reduce organic and nutrient content, enabling efficient downstream water treatment by conventional municipal water treatment technologies.

While studies have investigated in detail the microbial communities inhabiting fractured subsurfaces (6-8) and shifts in their structures over time in FFBs (9-11), there exists minimal information on the stability of these communities in ambient, surface, or storage conditions over time. Foci of applied research into the microbiology of PW systems includes considerations of pipeline corrosion via microbial-induced corrosion (MIC) (12-14), biocides for the prevention of well souring (15), and in-lab engineered microbial mats for treatment of PW (16). However, the effectiveness of simple chemical and/or microbial community additions to aerated PW for treatment improvement remains under-examined.

Herein, we have investigated the possibility of treating PW with native microbial communities through two lines of inquiry: 1) what chemical additions increase microbial treatment of produced water under aerated conditions?; and 2) what microbial communities are correlated with achieved improvements? These questions were investigated in a simplified setting whereby PW was stored at ambient temperatures in 1 m^3^ plastic totes for weeks at a time before addition into our experiment—mimicking the realities of PW treatment plans on-site at O&G operations. Here, we empirically determined changes in PW microbial ecology during storage and probe the subsequent behavior of those communities in an aerated treatment system. The ability for microbial communities native to PW to ‘self-treat’ water (i.e., improve water quality by the activity of native microbes alone) when mechanically aerated are not understood. Further, amendments of either KH_2_PO_4_ or AS test whether or not treatment of PW by native and / or introduced microbial communities can be improved. A better understanding of the biology underpinning successful treatment schemes is necessary for process upscaling and next-generation technology development—both of which are informed by this investigation. Lessons-learned about microbial ecology and the energetics thereof in the deep hydraulically fractured subsurface (6, 7) are yet to be optimized in treatment technologies. Subsurface-derived microorganisms possibly responsible for self-treatment of PW in aerated systems inherently require carbon, nitrogen, and other key elements (such as phosphorous) for assimilation into biomass. Therefore, the growth of microorganisms can remove undesirable constituents from water—suggesting that pathways to water treatment optimization by native microbial communities may exist.

Experiments were conducted to assess the viability of PW treatment with mechanical aeration as the primary mechanism to facilitate biological treatment. Microorganisms (primarily bacteria and archaea) in PW are of measurable abundance and, if aerated, may possess metabolisms capable of lowering concentrations of undesirable organic and inorganic constituents in PW—either alone or with the assistance of chemical and biological amendments. Through a controlled, long-term water treatment experiment we paired water chemistry and treatment data with 16S rRNA gene taxonomic profiling to investigate biological mechanisms of PW treatment efficacy—both under PW alone (baseline) and amendment conditions. This study represents a pairing of biological and chemical data throughout a simple analysis of the potential for self-treatment of PW in comparison to improved treatment by two widely available amendments (either KH_2_PO_4_ or AS). PW is known to be phosphate-limited for microbial growth and activity (4), suggesting that simple amendments of either phosphate (as KH_2_PO_4_) or microbial communities and nutrients (including phosphorous) from a traditional AS source may affect microbial consortia and their resultant treatment capabilities on PW. Herein, we experimentally verify the role of microbial consortia in PW treatment; further, we expound on ecology-informed alterations to PW treatment engineered systems that may optimize PW treatment under realistic, ambient settings—either in the lab or on-site at O&G operations.

## 2. Materials and methods

### 2.1. Produced water sourcing

PW was acquired approximately every 3 weeks from the Denver-Julesburg (D-J) basin in Weld County, Colorado. The water was collected from the same well pad throughout the duration of the study. The PW was stored in the laboratory in a closed, 1 m^3^ plastic tote at indoor ambient temperature (20 °C); this untreated PW was used as the influent (hereafter referred to as ‘IN’) for all reactors.

### 2.2. Reactor experiment

Three 2-L reactor vessels filled with PW were operated constantly with a repeated one-hour cycle: mixing and aeration for 30 minutes, and then mixing without aeration for 30 minutes to simulate a sequencing batch reactor (SBR). The system was operated at ambient lab temperatures (i.e., 20 °C) over the course of 256 days. One reactor was fed with only PW and was used as a control (baseline), while two additional reactors were used to test the effect of specific additives on both microbial communities and treatment efficiency (i.e., organic and nutrients removal).

Every 24 hours, aeration and mixing were paused, solids were allowed to settle for at least 20 minutes, and then 600 mL of liquid was decanted from each vessel. 600 mL of raw PW was then added in order to maintain a 2 L volume in each reactor. At this point, one experimental reactor was dosed with 200 mg of return activated sludge (RAS) from a municipal sequence batch membrane bioreactor (SB-MBR) (5), and the second reactor was dosed with 7.5 mg/L of phosphorous as KH_2_PO_4_. Due to evaporation, deionized water was added to each jar every 3-4 days to maintain the complete 2 L volume.

The 600 mL of decanted supernatant was used for water chemistry analyses. On study days 80 and 221, 45 mL of mixed liquor was collected prior to pausing the aeration cycle, followed by a 550 mL decant after settling. The 45 mL samples of mixed liquor were frozen immediately at –20 °C in 50 mL sterile polypropylene conical tubes (Corning Science Mexico, Reynosa, Tamaulipas, Mexico) until downstream microbiological processing. On study days 39, 80, and 102, 45 mL of raw produced water influent (IN) was collected, frozen, and stored with the same protocol as for reactor microbiome samples.

### 2.3. Chemical analysis

Anion measurements were conducted via ion chromatography (IC) on a ThermoFisher Dionex ICS-900, including a ThermoScientific DS5 conductivity detector, a Dionex ACRS 500 2 mm suppressor, a Dionex IonPac AG14A-5 um RFIC 3×30 mm guard column, and a Dionex IonPac AS14A-5 um RFIC 3×150 mm column. Metal analyses were conducted by inductively coupled plasma atomic emission spectroscopy (ICP-AES) on a Perkin Elmer Optima Model 5300 dual view spectrometer.

pH, conductivity, and turbidity were measured on daily calibrated benchtop instruments. sCOD, COD, ammonia, and ortho-phosphate were measured using Hach^®^ TNT PlusTM (Loveland, CO) reagent vials and Hach DR 6000TM spectrophotometer. Total suspended solids, fixed suspended solids, and volatile suspended solids were measured following Standard Method 2540.

### 2.4. DNA extraction

The periodically collected 45 mL mixed liquor samples from IN and reactors were frozen in 50 mL polypropylene conical tubes that were later thawed over 30 minutes at the time of microbiological processing. 40 mL of each sample (samples normalized to 40 mL for centrifuge balancing) were then centrifuged at 7,000 G for 45 minutes. The supernatant was decanted and the solid pellet was frozen until DNA extraction kit procedures. The ZymoBIOMICS DNA MiniKit was used according to manufacturer instructions to extract DNA from each pellet. Prior to DNA extraction, each pellet was thawed over 5 minutes; the lysis solution from the Zymo kit was then added to each sample conical. Conicals were vortexed until the pellet was completely solubilized in the lysis solution. The solution/pellet mixture was added to Zymo bead beating tubes, and the remainder of the DNA extraction protocol was conducted according to manufacturer instructions.

### 2.5. DNA sequencing

DNA amplicons were shipped to the Duke Center for Genomic and Computational Biology and sequenced on an Illumina MiSeq (Illumina Inc., San Diego, CA) instrument with V2 PE250 methods. 16S rRNA genes were amplified by PCR in duplicate 25 µL reactions. Reaction concentrations: 1x QuantaBio 5PRIME HotMasterMix, 0.2 µM 515-Y Forward Primer (5’-**GTA AAA CGA CGG CCA GT** CCG TGY CAG CMG CCG CGG TAA-3’) where the M13 forward primer (bold) is ligated to the 16S RNA specific sequence with a ‘CC’ spacer connection, 0.2 µM 926R (5’-CCG YCA ATT YMT TTR AGT TT-3’) Reverse Primer, and sample DNA. Primers are from Parada et al. (2015) (17) with the modification of forward primers by the addition of the M13 linker sequence (18). PCR was carried out on a Techne TC-5000 thermocyler. PCR parameters: primary denaturation at 98 °C for 30 s, followed by 30 cycles of 98 °C for 10 s, then 52 °C for 20 s and 72 °C for 10 s, followed by a final sequence extension of 72 °C for 5 min and a 10 °C hold. Six additional cycles of PCR with the same parameters as above were then performed to attach barcodes (19) to the M13 linker region of the forward primer (18). 16S rRNA amplicon cleanup was performed with KAPA Pure Beads (Kapa Biosystems, Indianapolis, IN) at 0.8x concentration according to the manufacturer’s instructions.

### 2.6. Bioinformatics

DADA2 (20) was used to—in order—remove forward and reverse primers from sequences; determine true sequence variants; merge paired-end reads; build a sequence table; and detect / eliminate chimeras. Untreated influent PW (IN) samples were rarefied to a sequencing depth of 12,874 sequences; i.e., all IN samples were forced to have the same sequencing depth via random subsampling. All IN samples had a sequencing depth greater than or equal to 12,874 and could be rarefied without rejection due to shallow sequencing depth. All experimental and control reactors were rarefied to 6,018 sequences and all samples from these vessels, similarly, were retained for post-rarefaction downstream analyses. Both rarefaction levels were based on an analysis of rarefaction curves, and values were chosen that allowed inclusion of all samples without substantial sacrifice of alpha diversity. For reproducibility of our code, a specific random start seed was defined for sequence rarefaction. Taxonomy assignments were called with a greengenes training file (version 13.8) (21) through the DADA2 taxonomy assignment script. Chloroplast and mitochondrial sequences were removed such that only bacterial and archaeal sequences remained (the greengenes database does not include eukaryotes). Operational taxonomic unit (OTU) tables, taxonomy tables, and sample meta-data were compiled within the R package ‘phyloseq’ (22).

## 3. Results

### 3.1. Raw PW and experimental reactor chemistry

Water quality analysis by mean variables summarized in Table 1 was conducted on untreated influent PW (IN) and reactors daily or weekly (sampling frequency is summarized in a table in our raw chemistry data submission), just prior to refilling of all 2 L vessels (after decanting/sampling). Throughout the study, samples from IN and all reactors were taken for IC (major anion) and ICP-AES (metal) analyses; mean values over the course of the study and the number of measurements (‘n’) are summarized in Table 2. Both AS and KH_2_PO_4_ additions to reactors improved ammonia, sCOD, and COD removal relative to the no-addition baseline throughout the duration of the study (Fig. 1, Table 1).

**Table 1.** Mean values of chemistry measurements throughout the study (plus or minus one standard deviation), and % removal (by mean) of same variables from IN levels by reactor addition type. Data are adapted from Nicholas, E.R. 2017 (23, 24). ‘n’—the number of measurements for each variable—are reported in a table in our raw chemistry data submission. ‘n.m.’ indicates not measured.

| | Raw PW<br>(IN) | Baseline | Activated<br>Sludge<br>(AS) | $\text{KH}_2\text{PO}_4$ | Mean %<br>removal by<br>Baseline | Mean %<br>removal by<br>AS | Mean %<br>removal by<br>$\text{KH}_2\text{PO}_4$ |
| --- | --- | --- | --- | --- | --- | --- | --- |
| pH | 6.84±0.32 | 8.08±0.14 | 8.08±0.24 | 8.05±0.29 | n.a. | n.a. | n.a. |
| Turbidity (NTU) | 89.9±73.6 | 82.3±63.4 | 832±624 | 348±289 | 8.45 | -826 | -287 |
| TSS (mg/L) | 43.8±46.2 | 138±257 | 1193±1233 | 292±326 | -215 | -2621 | -566 |
| VSS (mg/L) | 20.7±21.5 | 66±182 | 847±503 | 146±143 | -220 | -3992 | -604 |
| sCOD (mg/L) | 1509±859 | 1093±169 | 717±195 | 754±215 | 27.6 | 52.5 | 50.0 |
| COD (mg/L) | 1564±967 | 1102±300 | 1067±355 | 861±188 | 29.5 | 31.8 | 45.0 |
| Ammonia (mg/L) | 24.9±2.5 | 20.5±5.3 | 10.2±3.3 | 9.6±2.9 | 17.8 | 59.2 | 61.5 |
| Phosphate (mg/L) | n.m. | 0.07±0.13 | 0.85±2.46 | 16.8±28.2 | n.m. | n.m. | n.m. |
| Conductivity (mS/cm) | n.m. | 28.9±4.6 | 28.3±3.9 | 29.2±4.0 | n.m. | n.m. | n.m. |

**Table 2.** Mean concentrations of metals (mg/L) (for IN, n = 3; for reactors, n = 4) and anions (mg/L, except for IN which is in ppm) (for IN, n = 4; for reactors, n = 5), throughout the study. Metals and anions were determined via ICP-AES and IC, respectively. ‘n.a.’ indicates below instrument detection limit and ‘n.m.’ indicates not measured.

| Metal / Anion | Raw PW (IN) | Baseline | Activated<br>Sludge (AS) | KH<sub>2</sub>PO<sub>4</sub> |
| --- | --- | --- | --- | --- |
| Al | n.a. | n.a. | n.a. | n.a. |
| As | 0.12 | 0.41 | 0.24 | n.a. |
| B | 16.7 | 20.1 | 16.5 | 17.6 |
| Ba | 9.07 | 10.5 | 8.47 | 7.55 |
| Be | n.a. | n.a. | n.a. | n.a. |
| Ca | 308.8 | 341.9 | 298.2 | 306.4 |
| Cd | 0.00 | 0.02 | 0.01 | n.a. |
| Co | n.a. | n.a. | 0.02 | 0.02 |
| Cr | 0.01 | 0.03 | n.a. | 0.03 |
| Cu | 0.04 | 0.22 | 0.12 | 0.22 |
| Fe | 1.32 | 4.37 | 1.69 | 0.75 |
| K | 33.9 | 41.7 | 38.7 | 88.8 |
| Li | 5.02 | 5.46 | 4.71 | 5.26 |
| Mg | 35.0 | 40.7 | 37.9 | 37.5 |
| Mg | 34.8 | 29.2 | 28.9 | 27.8 |
| Mn | 0.45 | n.m. | n.m. | n.m. |
| Na | 5157 | 5511 | 5121 | 5506 |
| Ni | n.a. | 2.17 | 3.61 | 7.90 |
| P | n.m. | 1.13 | 3.11 | 10.8 |
| Pb | 0.11 | 9.27 | 7.67 | 5.33 |
| S | n.m. | 5.42 | 5.82 | 6.60 |
| Se | n.a. | 13.8 | 10.3 | 31.3 |
| Si | 22.0 | 34.3 | 31.1 | 30.6 |
| Sr | 38.4 | 39.9 | 36.2 | 37.7 |
| Tl | 0.05 | 0.10 | n.a. | n.a. |
| V | n.a. | 0.47 | 0.31 | 0.23 |
| Zn | n.m. | 0.37 | 0.32 | 0.32 |
| Sn | n.a. | n.a. | n.a. | n.a. |
| Mo | n.a. | n.a. | n.a. | n.a. |
| Sb | n.a. | n.a. | n.a. | n.a. |
| Ti | n.a. | n.a. | n.a. | n.a. |
| Fluoride | n.a. | n.a. | n.a. | n.a. |
| Chloride | 8807 | 9703 | 8404 | 9296.1 |
| Nitrite | 6.06 | 12.2 | 9.88 | 12.7 |
| Bromide | 98.6 | 106.4 | 125.8 | 104.6 |
| Nitrate | n.a. | 11.1 | n.a. | 16.6 |
| Phosphate | n.a. | n.a. | n.a. | 99.4 |
| Sulfate | 5.9 | 24.21 | 14 | 37.6 |

**Fig. 1.**
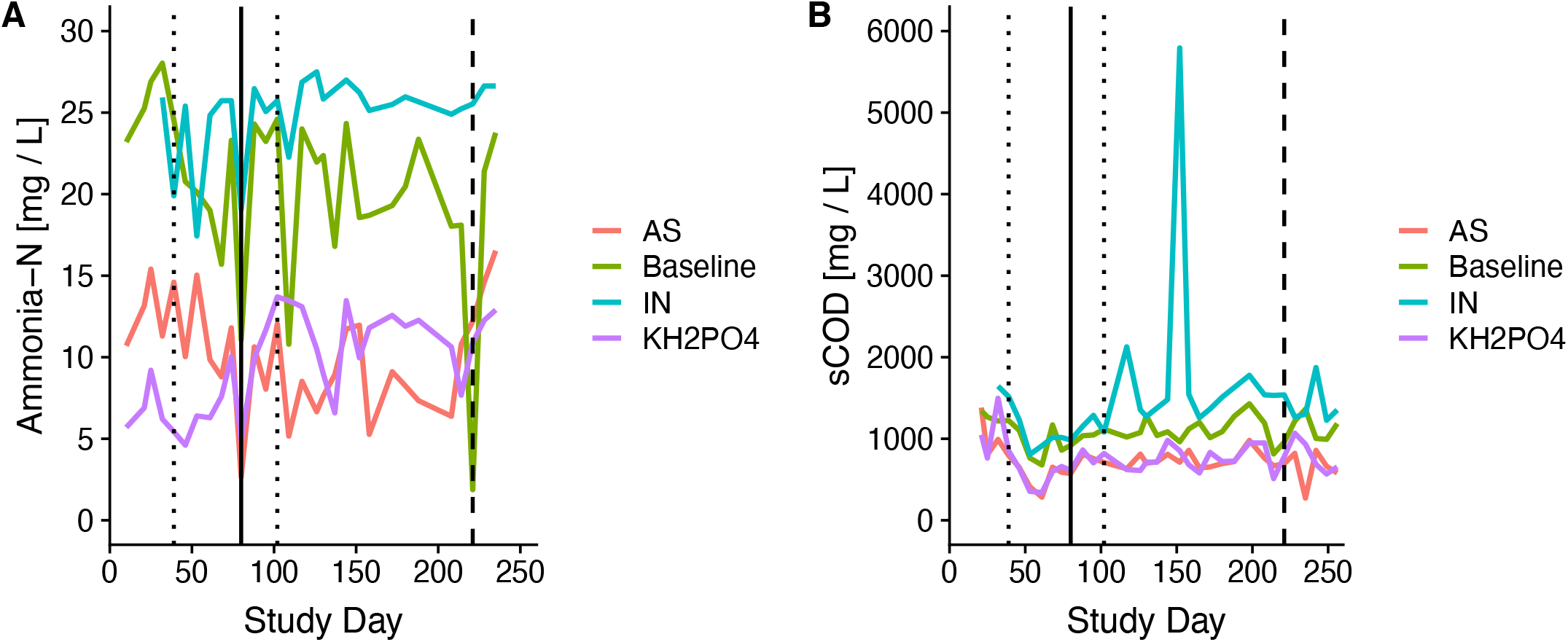
Concentrations of ammonia-N (**A**) and sCOD (**B**) in IN, and reactors (baseline, AS and KH_2_PO_4_). Vertical dotted lines indicate the time points of biological sampling of IN; the vertical dashed line indicates a time point of biological sampling of reactors; the vertical solid line indicates biological sampling of both IN and reactors.

### 3.2. Raw PW influent (IN) microbial community

IN fed into the reactor experiments was analyzed for order taxonomy microbial abundance over time (Fig. 2). During storage times of roughly 3 weeks in 1 m^3^ plastic totes, IN (measured at time of feed into reactors) coalesced into just a handful of taxonomic orders with a relative abundance greater than 1%. Similarly, the number of observed operational taxonomic units (OTUs) in IN decreased with time (Table 4).

**Table 4.** Operational taxonomic unit (OTU) alpha diversities and cumulative rate of uncommon taxonomic classifications throughout the study.

| Untreated Influent or Reactor Type | Study day | # Observed OTUs | % Bacteria not in top 10 orders ('Other Bacteria') |
| --- | --- | --- | --- |
| IN | 39 | 135 | 17.8 |
|  | 80 | 75 | 18.2 |
|  | 102 | 75 | 15.5 |
| Baseline | 80 | 80 | 30.1 |
|  | 221 | 55 | 7.2 |
| AS addition | 80 | 113 | 39.5 |
|  | 221 | 329 | 43.9 |
| KH<sub>2</sub>PO<sub>4</sub> addition | 80 | 28 | 2.2 |
|  | 221 | 38 | 7.0 |

**Figure 2.**
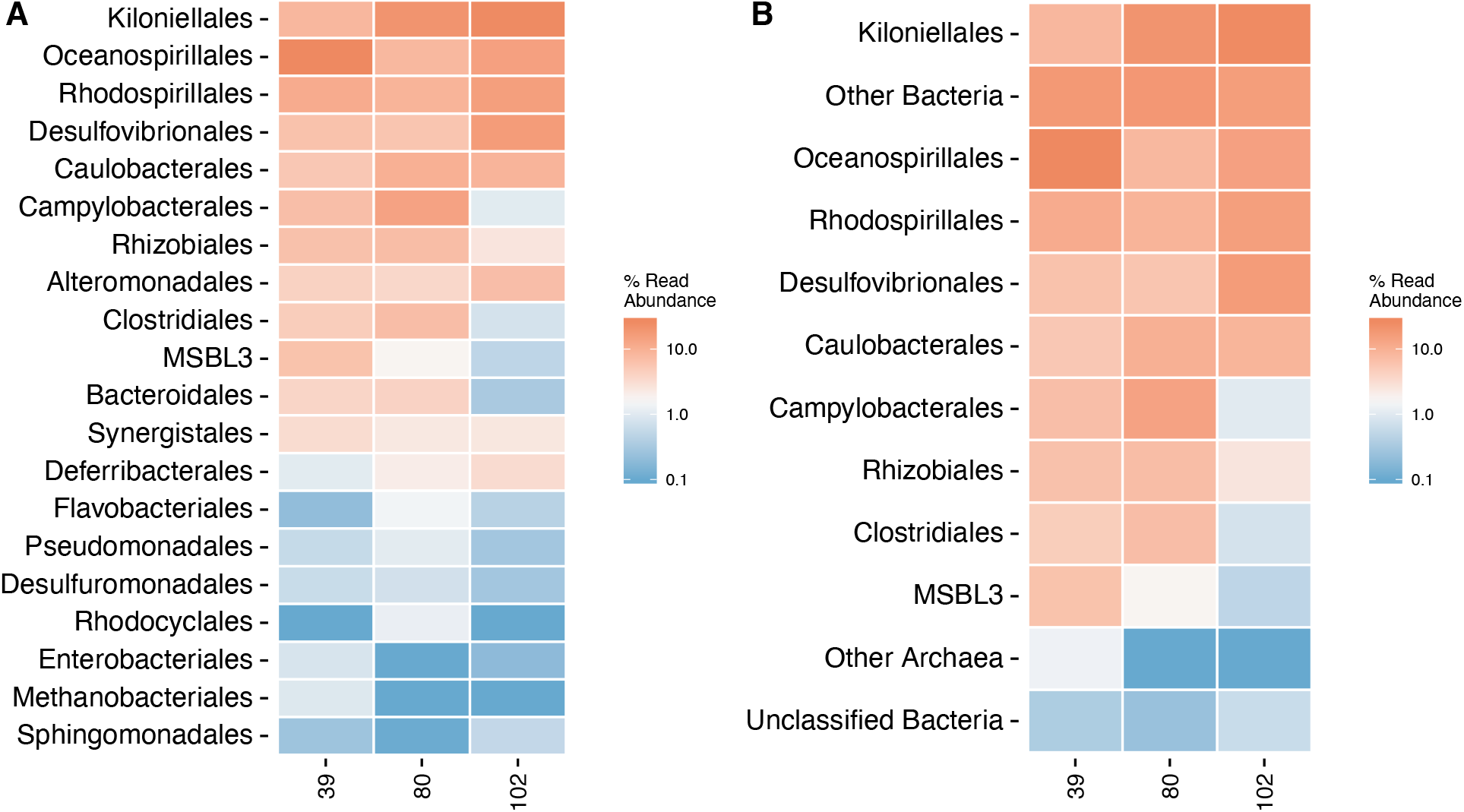
IN order taxonomy relative abundances vs. day in study. The top 20 most abundant orders **(A)** are contrasted with a modified chart **(B)** where bacteria that are not one of the top 10 most abundant orders are grouped as ‘Other Bacteria’; i.e., all orders of bacteria that are not one of the 10 most common—but have known order taxonomy—are compiled into a single, shared, category denoted ‘Other Bacteria’. ‘Other Bacteria’, when treated as a composite group, become the second most common taxonomic classification (panel **B**). Bacteria that could not be classified are plotted separately as ‘Unclassified Bacteria’.

Genus *Thalassospira* ranked as both the most prevalent and persistent genera of bacteria over time in IN; its relative abundance within classified genera increased from 11.1% to 22.4% and then to 28.4% over time. In contrast, genus *Marinobacterium* (the second most common genera in IN) decreased in abundance over time, beginning at 34.8% before decreasing to 8.3% and then slightly increasing to 15.9%.

### 3.3. Experimental/baseline reactor microbial community response

Both the baseline- and KH_2_PO_4_- amended reactor vessels consisted almost exclusively of phylum *Proteobacteria*, while the AS vessel had greater diversity at the phylum level across all time points—the AS reactor contained dominant phyla Proteobacteria (62.1 - 62.2%), Bacteroidetes (11.5 - 28.7%), and Chloroflexi (1.6 – 9.7%). To more closely assess biotic differences between vessels, the order taxonomic rank was chosen to show the relative abundances of taxa in each reactor (Fig. 3). Order taxonomy was chosen because the vast majority of sequences from both IN and reactor samples were identifiable at this level. Furthermore, order taxonomy provided a greater resolution of differences between samples than the phylum level. At both reactor biological sampling time points (day 80 and day 221), all experimental vessels were distinct from one another as well as from the baseline reactor (Fig. 3). The baseline reactor had similar compositions on day 80 and day 221, and the relative abundances of present taxa varied only slightly, indicating dominant-taxa stability (Fig. 3). Specifically in the baseline vessel, orders *Kiloniellales* and *Caulobacterales* were most common and persisted over time between days 80 and 221; this is in contrast to IN in which *Kiloniellales* and *Oceanospirillales* were most abundant. Reactors dosed with KH_2_PO_4_ appear laden with completely different taxonomic orders relative to both the baseline and AS reactors, the most common of which were *Rhodobacterales, Pseudomonadales*, and *Oceanospirillales* (Fig. 3). While some orders have a noticeable presence in both the baseline vessel and the vessel dosed with KH_2_PO_4_, the relative abundances of these shared taxa were remarkably different. Stark differences in community structure between the AS and KH_2_PO_4_ addition reactors are orthogonal to their similar removal efficacy (Fig. 1).

**Figure 3.**
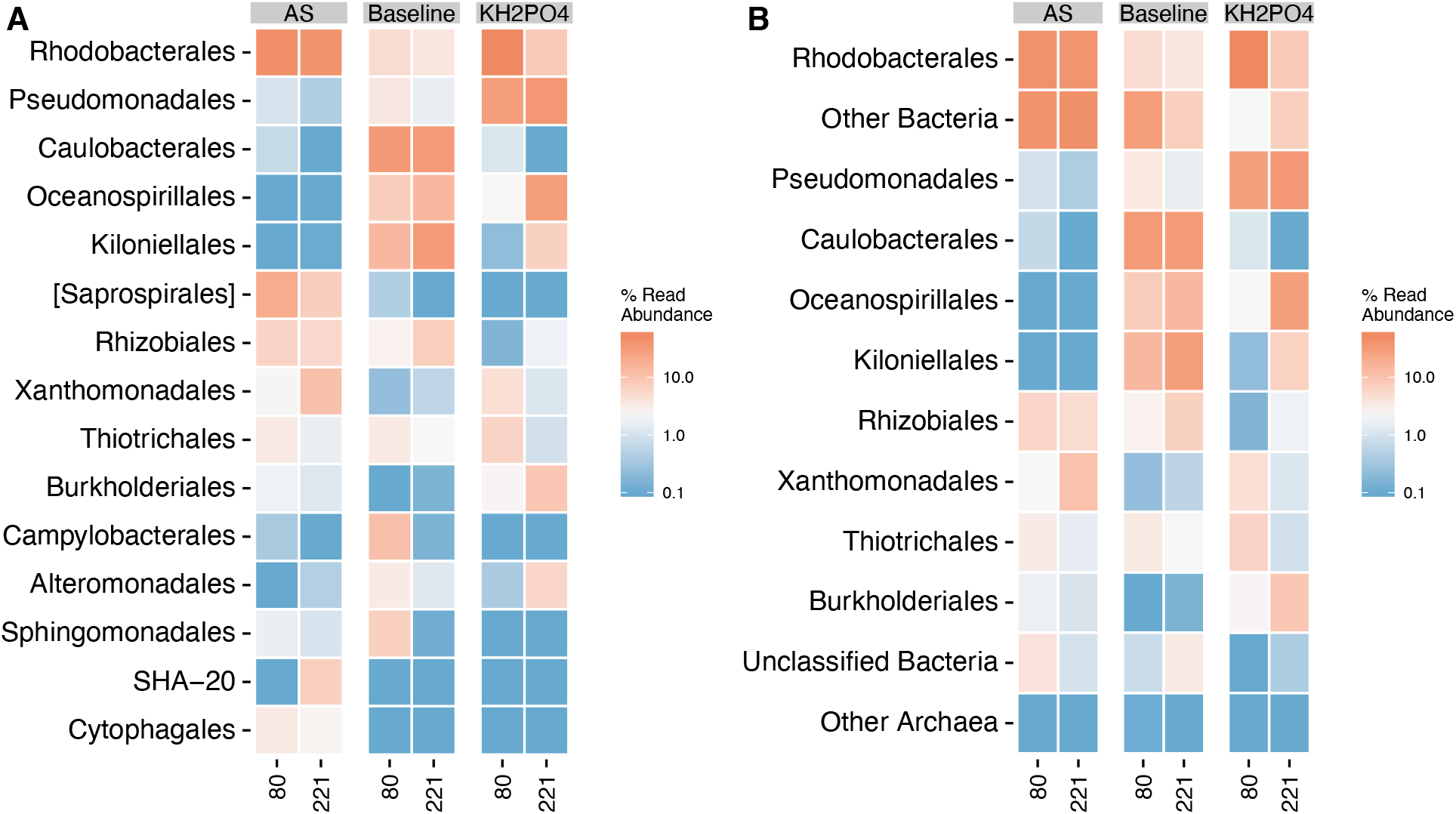
Order taxonomy relative abundances vs. day in study for AS and KH_2_PO_4_ amendment reactors compared to baseline **(A)**. Panel **A** is modified in panel **B** by combining orders of bacteria that are not one of 10 most common into a new group called ‘Other Bacteria’ **(B)**; i.e., all orders of bacteria that are not one of the 10 most common—but have known order taxonomy—are compiled into a single, shared, category denoted ‘Other Bacteria’. ‘Other Bacteria’, when treated as a composite group, become the second most common taxonomic classification (panel **B**). Bacteria that could not be classified are plotted separately as ‘Unclassified Bacteria’.

Notably, at the genus taxonomic rank, *Thalassospira* existed at no more than 7.7% relative abundance in either the AS or KH_2_PO_4_ reactors at both time points, marking a substantial reduction in abundance from IN. However, genus *Marinobacterium* (the second most common genera in IN) completely disappeared from the AS vessel but remained highly abundant in the KH_2_PO_4_ vessel. In the baseline reactor, the most common genera were largely unchanged from IN: *Thalassospira, Marinobacterium*, and *Mycoplana* remained the most abundant.

### 3.4. Microbial Diversity Response to Experiments

As time progressed microbial community diversity increased within both the AS and KH_2_PO_4_ test groups while the diversity in the baseline reactor decreased with time (Table 4). ‘Other Bacteria’, existing at nominally low relative abundances as individuals—but have known order taxonomy, collectively represented a substantial component of the total community in all of baseline, AS, and KH_2_PO_4_ conditions (Fig. 3B, Table 4). The baseline, AS, and KH_2_PO_4_ reactors showed different trends in the percent of identified bacteria not in the top 10 most abundant orders (i.e., ‘Other Bacteria’). The AS condition had an abundant coalition of less common microbes (i.e., microbial community members in low abundance representative of the so-called rare biosphere); though less pronounced, these lesser-appreciated constituents of a canonical ‘abundant’ microbiome were still a substantial component of both the baseline and KH_2_PO_4_ reactors (Table 4).

## 4. Discussion

Results demonstrate that additions of AS or KH_2_PO_4_ were simple and effective means to enhance the pretreatment of PW in an intermittent aeration/mixing system. Pretreatment of PW in a microbial system was viable over the extended study duration, even when PW was stored for weeks at indoor ambient lab conditions. Periodic peaks in PW feed water quality (e.g., sCOD (Fig. 1B)) were well tolerated by the system, suggesting that pretreatment of PW can be achieved on-site at O&G operations with little intervention. Here, we report that ammonia, sCOD, and COD removal was modest with intermittent aeration cycles in the baseline reactor while removal of the same chemical constituents was substantially enhanced by the addition of either AS or KH_2_PO_4_. Given that the raw PW (IN) fed into the experimental and baseline reactors had little-to-no detectable phosphate (Tables 1 and 2), and the subsequent measurable concentrations of phosphate during treatment (Tables 1 and 2), it is likely that the microbial consortia contained within our treatment scheme were nutrient limited with respect to phosphorous prior to AS or KH_2_PO_4_ addition.

The KH_2_PO_4_ reactor had lower microbial diversity relative to the baseline reactor, but had better treatment performance; however, we noticed that the reactor with added AS had a similar increase in treatment performance over baseline, but with a complete opposite microbial diversity response—i.e., the number of observed OTUs was 41% greater relative to the baseline reactor on day 80, and is 498% greater on day 221 (Table 4). The changes in relative abundances of the most common order-classified taxa are deceivingly slight in comparison to shifts in the composite abundance of less common microbiota—all with known order taxonomy; in both the raw PW feed and the experimental reactors, greater changes in ‘Other Bacteria’ and the number of observed OTUs are seen (Table 4) relative to what may be perceived were one to look only at the top-most abundant orders. Collectively, comparisons of the most abundant taxa, the number of observed OTUs, and the contribution of uncommon microbiota to the total population suggest that a broad interpretation of ‘who’s there’ is essential for a thorough assessment of biological PW treatment systems.

As observed in our experimental setups, it may be well true that microbial community composition of the rarer members of the community, those in least abundance and possibly present as mere singletons, likely play a substantial role in both community support to abundant members and functional redundancy of metabolic response (25, 26). Microbiota native to PW and those that live within AS are not identical. Nonetheless, we empirically show functional redundancy between these two distinct communities; i.e., similar PW treatment efficacy is conducted by both. Louca et al. (25) describe community functional capacity as decoupled from the abundance of taxonomic clades; further, this decoupling is thought to systemically arise from the need for microbial communities to possess a base set of critical metabolic functions. In our experiment, it is possible that stochastic processes are not responsible for the similar treatment efficacy of KH_2_PO_4_- and AS-additions but, rather, these critical metabolic functions are actively selected for from distinctly different microbial communities; i.e., an inherent property of microbial communities native to PW in engineered systems may be to adapt to the task at hand independent of the overwhelming diversity of microorganisms within ‘The deep, hot biosphere’, as hypothesized by Thomas Gold in 1992 (27, 28).

Genus *Thalassospira* was most common in the raw PW feed and is a known chemotactic bacterium that pursues inorganic phosphate (29), further suggesting that raw PWs are initially phosphorous-limited. In addition to genus *Thalassospira*, genus *Marinobacterium* (a known carbon opportunist in O&G fracture systems (7)) was found to be persistent over time in the raw PW feed, despite a reduction in alpha diversity as time progressed (Table 4); the persistence of microbiota native to subsurface systems in PW suggests that the functional capacity to treat PW may prevail under ambient, surface conditions. Persistence of subsurface metabolisms in surface, engineered treatment systems may enable more techno-economic flexibility in the treatment of PW for beneficial reuse. Genus *Marinobacterium* was also common in the baseline and KH_2_PO_4_-amended reactors; however, despite similar chemistry removal performance of the AS-amended reactor, *Marinobacterium* in those samples was neither present nor enriched with time. Disparate fates of *Marinobacterium* in the experimental reactors suggest that there exist multiple microbiological consortia capable of treating PW, and that some of those communities—such as those inoculated by AS—may not necessarily derive exclusively from the subsurface or PW.

In light of Frank, et al. (4), it is likely that the treatment of PW in POTWs—seen optimally between 6-20% by volume—is a synthesis of performance by both AS and native PW microbial communities. Nonetheless, microbial communities existing within AS may have a limit on their ability to treat PW—suggesting that simple additions of phosphorous—optimized within a range centered at daily doses of 7.5 mg P/L—in an aerated/mixed system could be an excellent mechanism to enhance pre-treatment of PW. Further, entirely different microbial consortia appear to have similar chemistry removal efficacy; this is promising for the treatment of PW because an exact replication of a particular treatment-capable microbial consortia may not be necessary, indeed likely impossible given the variance of PW microbial communities, to reach treatment goals.

At the order taxonomy level, *Rhodobacterales* are the most greatly enriched taxa in both AS- and KH_2_PO_4_-amended reactors across all time points (Fig. 3). This increase in relative abundance may indicate a role in PW treatment by *Rhodobacterales. Rhodobacterales* are known surface colonizers in saline coastal marine waters (30); their exceptional adaptation for early surface colonization may position them well for establishment as a biofilm in PW treatment reactors. Shortly after exposure to crucial nutrient additions (i.e., phosphorous), it is possible that biofilm colonization by *Rhodobacterales* is potentiated—which would make sense in light of *Rhodobacterales*’s enrichment in AS- and KH_2_PO_4_-amended reactors. Future work should consider the role of planktonic versus biofilm communities in the treatment of PW, revealing more optimal physical designs of treatment reactors better tuned to the needs of surface colonizers.

## 5. Conclusion

Herein, we have described a simple, minimal-intervention bench scale PW treatment scheme. Microbial consortia native to PW are capable of attenuating concentrations of sCOD, COD, and ammonia. Further, simple additions of either AS or phosphorous as KH_2_PO_4_ are sufficient to further improve water treatment. Microbiome analyses of this simple, O&G field-site compatible engineered system suggest that several microbial consortia can treat PW across a range of both chemical and environmental conditions—a promising finding for practical considerations of PW treatment system implementation in the field at O&G sites where advanced systems are less tractable and it is essential to consider techno-economic viability. Via simple additions of AS or KH_2_PO_4_, PW (a phosphorous-limited system) water quality can be enhanced by microbial communities jumpstarted by these additions. In light of this study, future work on the treatment of PW should consider PW a ‘living system’. Both microbiota native to the subsurface and introduced to PW by AS apparently tolerate changes in environmental conditions while consistently enhancing the water quality of PW. Future work should focus on greater understanding of the mechanisms of ecological functional redundancy in engineered PW treatment systems; improved knowledge about basic ecological phenomena during applied interventions will likely lead to better optimized PW treatment schemes for beneficial water reuse.

## Data Availability

All raw data, R code, files, and R objects used in the production of this manuscript are available for review and enable immediate reproduction of results (link to DropBox below).

https://www.dropbox.com/sh/7fpfkc0lu3d28sa/AACsyRAeWTXx773BK1SPSIEma?dl=0

If the manuscript is accepted, all raw DNA sequencing reads will be uploaded to NCBI SRA with accession numbers listed here; additionally, the same raw data folder shared for review on DropBox will be shared publicly on figshare with a DOI listed here.

## Acknowledgements

Financial support for this work was provided by ConocoPhillips through their contribution to the Center for a Sustainable WE^2^ST (Water-Energy Education, Science, and Technology) at the Colorado School of Mines. The authors also acknowledge the support provided by the NSF/SRN program under Cooperative Agreement No. CBET 1240584, the Edna Bailey Sussman Foundation Internship, and the National Science Foundation Graduate Research Fellowship (Fellow I.D. No. 2019258966). The authors also thank Maria L. Day, Estefani Bustos, Mike Veres, and Tani Cath for their assistance with laboratory analyses and technical support.

